# Visual experience shapes the neural networks remapping touch into external space

**DOI:** 10.1101/149021

**Authors:** Virginie Crollen, Latifa Lazzouni, Mohamed Rezk, Antoine Bellemare, Franco Lepore, Olivier Collignon

## Abstract

Localizing touch relies on the activation of skin-based and externally defined spatial frames of references. Psychophysical studies have demonstrated that early visual deprivation prevents the automatic remapping of touch into external space. We used fMRI to characterize how visual experience impacts on the brain circuits dedicated to the spatial processing of touch. Sighted and congenitally blind humans (male and female) performed a tactile temporal order judgment (TOJ) task, either with the hands uncrossed or crossed over the body midline. Behavioral data confirmed that crossing the hands has a detrimental effect on TOJ judgments in sighted but not in blind. Crucially, the crossed hand posture elicited more activity in a fronto-parietal network in the sighted group only. Psychophysiological interaction analysis revealed that the congenitally blind showed enhanced functional connectivity between parietal and frontal regions in the crossed versus uncrossed hand postures. Our results demonstrate that visual experience scaffolds the neural implementation of touch perception.

**Significance statement:** Although we seamlessly localize tactile events in our daily life, it is not a trivial operation because the hands move constantly within the peripersonal space. To process touch correctly, the brain has therefore to take the current position of the limbs into account and remap them to their location in the external world. In sighted, parietal and premotor areas support this process. However, while visual experience has been suggested to support the implementation of the automatic external remapping of touch, no studies so far have investigated how early visual deprivation alters the brain network supporting touch localization. Examining this question is therefore crucial to conclusively determine the intrinsic role vision plays in scaffolding the neural implementation of touch perception.

## Introduction

Quickly and accurately localizing touch in space is crucial for efficient action planning toward an external stimulus making contact with the body. Although we seamlessly do it in daily life, it is not a trivial operation because the hands move constantly within the peripersonal space as different postures are adopted. Therefore, the brain must transform tactile coordinates from an initial skin-based representation to a representation that is defined by coordinates in external space (Yamamoto and Kitazawa, 2001; Shore et al., 2002; Azañón and Soto-Faraco, 2008; Azañón et al., 2010a, 2015). For example, when sighted individuals have to judge which of their two hands receive a tactile stimulation first (Temporal Order Judgment task – TOJ), they do much more errors when their hands are crossed over the body midline compared to when the hands are uncrossed (Yamamoto and Kitazawa, 2001; Shore et al., 2002; Heed and Azañón, 2014). This crossed-hands deficit has been attributed to the misalignment of anatomical and external frames of reference (Yamamoto and Kitazawa, 2001; Shore et al., 2002). Because the task requirements have nothing spatial (in theory, the task could be solved by using somatotopic coordinates only), this crossing-hand effect compellingly illustrates how the external remapping of touch is automatic in sighted people (Heed and Azañón, 2014). Specific brain networks including parietal and premotor areas have been demonstrated to support this automatic remapping of touch into an external spatial coordinate system (Lloyd et al., 2003; Matsumoto et al., 2004; Azañón et al., 2010a; Takahashi et al., 2013; Wada et al., 2012).

Congenitally blind people, in contrast, do not show any crossing-hand deficit when involved in a tactile TOJ task (Röder et al., 2004; Crollen et al., 2017). This suggests that the default remapping of passive touch into external spatial coordinates is acquired during development as a consequence of visual experience. Does the absence of visual experience also alter the neural network typically recruited when people experience a conflict between skin-based and external spatial coordinates of touch? Investigating how congenital blindness reorganizes the brain network supporting touch localization is crucial to conclusively determine the intrinsic role vision plays in scaffolding the neural implementation of the perception of touch location. In order to address this question, we used functional Magnetic Resonance Imaging (fMRI) to characterize the brain activity of congenitally blind individuals and sighted controls performing a tactile TOJ task with either their hands uncrossed or with the hands crossed over the body midline.

## Method

### Participants

Eleven sighted controls (SC) [four females, age range 22-64 y, (mean ± SD, 46 ± 14 y)] and 8 congenitally blinds (CB) participants [2 females, age range 24-63 y, (mean ± SD, 47 ± 13 y)] took part in the study (see Table 1 for a detailed description of the CB participants). The mean age of the SC and CB groups did not statistically differ (*t*(17) = 0.11, *p* = .92). At the time of testing, the participants in the blind group were totally blind or had only rudimentary sensitivity for brightness differences and no patterned vision. In all cases, blindness was attributed to peripheral deficits with no additional neurological problems. Procedures were approved by the Research Ethics Boards of the University of Montreal. Experiments were undertaken with the understanding and written consent of each participant. Both groups of participants were blindfolded when performing the task.

**Table 1.**
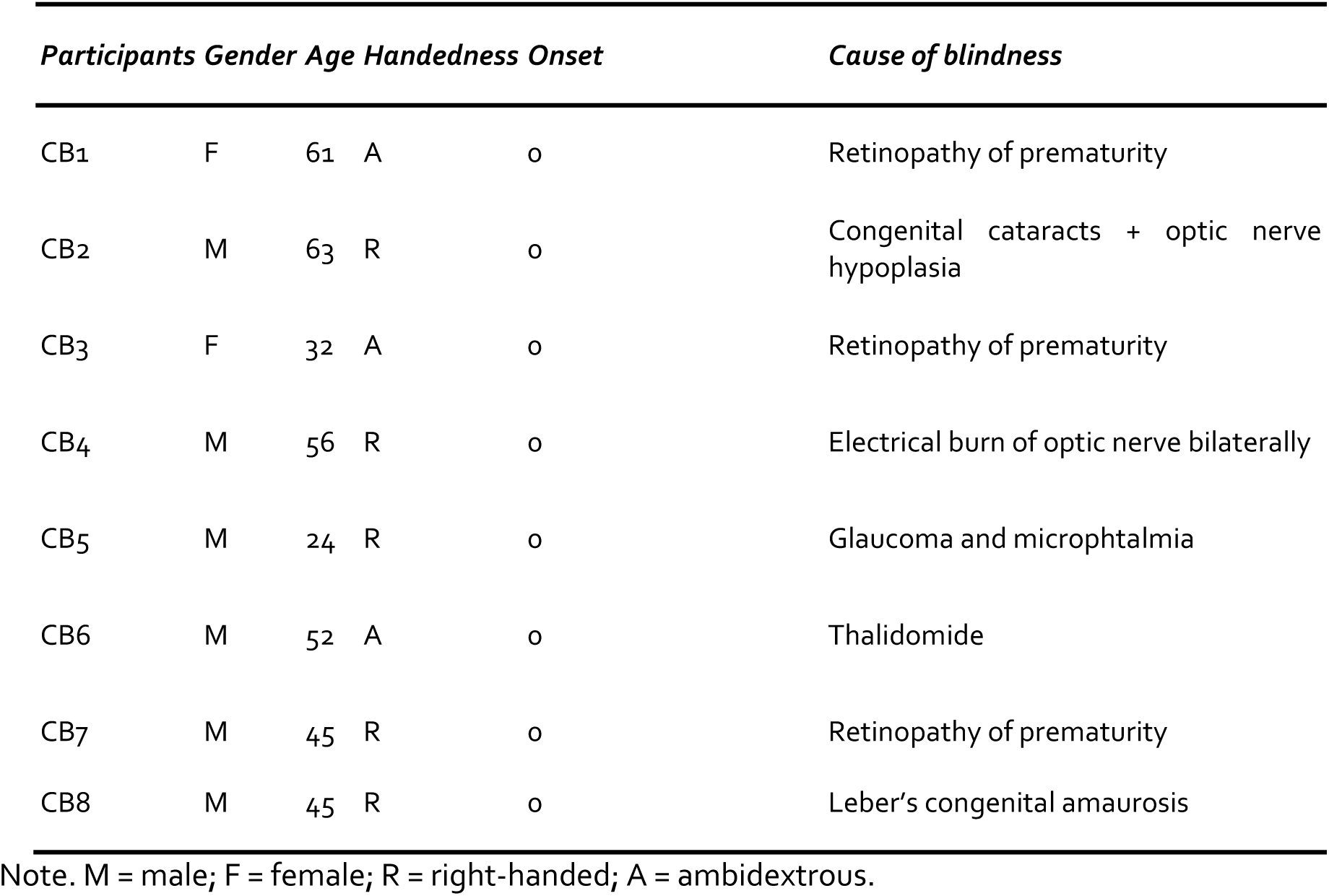
Characteristics of the blind participants

### Task and general experimental design

In this task, two successive tactile stimuli were presented for 50 ms to the left and right middle fingers at 6 different stimulus onset asynchronies (SOAs): -120, -90, -60, 60, 90, 120. Negative values indicated that the first stimulus was presented to the participant’s left hand; positive values indicated that the first stimulus was presented to the participant’s right hand. Tactile stimuli were delivered using a pneumatic tactile stimulator (Institute for Biomagnetism and Biosignal Analysis, University of Muenster, Germany). A plastic membrane (1 cm in diameter) was attached to the distal phalanxes of the left and right middle fingers and was inflated by a pulse of air pressure delivered through a rigid plastic tube. Participants had to press a response button placed below the index finger of the hand that they perceived to have been stimulated first. They had 3550 ms to respond otherwise the trial was terminated. Participants were asked to perform the task either with their hands in a parallel posture (i.e., uncrossed posture) or with their arms crossed over the body midline. Stimuli were delivered and responses were recorded using Presentation software (Neurobehavioral Systems Inc.) running on a Dell XPS computer using a Windows 7 operating system.

Participants were scanned in 2 fMRI sessions using a block design. One run consisted of 16 successive blocks (22 s duration each) separated by rest periods ranging from 11 to 14 s (median 12.5 s), during which participants had to perform the TOJ judgments either with the hands uncrossed or with the hands crossed. The starting run (uncrossed or crossed) was counterbalanced across participants. Each block, either uncrossed or crossed, consisted of 6 successive pairs of stimulations (each SOA was randomly presented once in each block).

Before the fMRI acquisition, all participants underwent a training session in a mock scanner, with recorded scanner noise played in the bore of the stimulator to familiarize them with the fMRI environment and to ensure that the participants understood the task.

### Behavioral data analyses

The mean percentages of “right hand first” responses were calculated for each participant, SOA and posture. These raw proportions were transformed into their standardized z-score equivalents and then used to calculate the best-fitting linear regression lines of each participant (Shore et al., 2002).

The just noticeable difference (JND; the smallest interval needed to reliably indicate temporal order) was secondly calculated from the mean slope data by subtracting the SOA needed to achieve 75% performance from the one needed to achieve 25% performance and dividing by 2 (Shore et al., 2002).This value was calculated for the entire group. It could not be determined independently for all observers because several sighted people obtained a slightly negative slope value for the crossed posture (Shore et al., 2002). This indicated that some participants responded with the opposite hand as the one that has been stimulated first (Yamamoto and Kitazawa, 2001).

### fMRI data acquisition and analyses

#### Acquisition

Functional MRI-series were acquired using a 3-T TRIO TIM system (Siemens, Erlangen, Germany), equipped with a 12-channel head coil. Multislice T2*-weighted fMRI images were obtained with a gradient echo-planar sequence using axial slice orientation (TR = 2200 ms, TE = 30 ms, FA = 90°, 35 transverse slices, 3.2 mm slice thickness, 0.8 mm inter-slice gap, FoV = 192×192 mm^2^, matrix size = 64×64×35, voxel size = 3×3×3.2 mm^3^). Slices were sequentially acquired along the z-axis in feet-to-head direction. The 4 initial scans were discarded to allow for steady state magnetization. Participants’ head was immobilized with the use of foam pads that applied pressure onto headphones. A structural T1-weigthed 3D MP-RAGE sequence (voxel size= 1x1x1.2 mm^3^; matrix size= 240x256; TR= 2300 ms, TE= 2.91 ms, TI= 900 ms, FoV= 256; 160 slices) was also acquired for all participants.

#### Analyses

Functional volumes from the uncrossed and crossed conditions were pre-processed and analyzed separately using SPM8 (<u>http://www.fil.ion.ucl.ac.uk/spm/software/spm8/</u>; Welcome Department of Imaging Neuroscience, London), implemented in MATLAB (MathWorks). Pre-processing included slice timing correction of the functional time series (Sladky et al., 2011), realignment of functional time series, co-registration of functional and anatomical data, a spatial normalization to an echo planar imaging template conforming to the Montreal Neurological institute space, and a spatial smoothing (Gaussian kernel, 8mm full-width at half-maximum, FWHM). Serial autocorrelation, assuming a first-order autoregressive model, was estimated using the pooled active voxels with a restricted maximum likelihood procedure and the estimates were used to whiten the data and design matrices.

Following pre-processing steps, the analysis of fMRI data, based on a mixed effects model, was conducted in two serial steps accounting respectively for fixed and random effects. For each subject, changes in brain regional responses were estimated through a general linear model including the responses to the 2 experimental conditions (uncrossed, crossed). These regressors consisted of a boxcar function convolved with the canonical hemodynamic response function. Movement parameters derived from realignment of the functional volumes (translations in x, y and z directions and rotations around x, y and z axes) and a constant vector were also included as covariates of no interest. We used a high-pass filter with a discrete cosine basis function and a cut-off period of 128s to remove artefactual low-frequency trends.

Linear contrasts tested the main effect of each condition ([Uncrossed], [Crossed]), the main effects of general involvement in a tactile TOJ task ([Uncrossed+Crossed]), the specific effect of the uncrossed condition ([Uncrossed >Crossed]) and the specific effect of the crossed condition [Crossed>Uncrossed]. These linear contrasts generated statistical parametric maps [SPM(T)]. The resulting contrast images were then further spatially smoothed (Gaussian kernel 8 mm FWHM) and entered in a second-level analysis, corresponding to a random effects model, accounting for inter-subject variance. One-sample t-tests were run on each group separately. Analyses characterized the main effect of each condition ([Uncrossed], [Crossed]), the main effect of general TOJ ([Uncrossed +Crossed]), the specific effects of the uncrossed ([Uncrossed >Crossed]) and the crossed condition [Crossed>Uncrossed]. Two-sample t-tests were then performed to compare these effects between groups ([Blind vs. Sighted]).

#### Statistical inferences

Statistical inferences were performed at a threshold of p < 0.05 after correction for multiple comparisons (Family Wise Error method) over either the entire brain volume, or over small spherical volumes (15 mm radius) located in structures of interest. Coordinates of interest for small volume corrections (SVCs) were selected from the literature examining brain activations related to the external representation of space in sighted participants.

### Standard stereotactic coordinates (x,y,z) used for SVC (in MNI space)

<u>*Frontal locations*</u> : Left precentral gyrus (preCG): -46, -10, 40 (Matsumoto et al., 2004); -46, 8, 46; 24, 4, 58 (Lloyd et al., 2003); -40, 4, 40 (Takahashi et al., 2013). Dorsolateral prefrontal cortex: -52, 14, 26 (Takahashi et al., 2013). *<u>Parietal locations:</u>* left precuneus (superior parietal lobule): -8, -56, 58; -14, -66, 52 (Matsumoto et al., 2004), -32, -54, 62 (Takahashi et al., 2013); right precuneus: 24, -44, 72 (Lloyd et al., 2003); right posterior parietal cortex (PPC): 26, -54, 42; 24, -54, 58 (Lloyd et al., 2003); 26, -58, 43 (Azañón et al., 2010); 24, -51, 42 (Zaehle et al., 2007); left PPC: -46, -64, 38 (Lloyd et al., 2003); left medial intraparietal area (MIP): -46, -52, 50 (Lloyd et al., 2003). Superior parietal gyrus: -26, -72, 32 (Takahashi et al., 2013). *<u>Temporal locations</u>*: right middle temporal gyrus : 46, -40, 2 (Takahashi et al., 2013).

#### Psychophysiological interaction

Psychophysiological interaction (PPI) analyses were computed to identify any brain regions showing a significant change in the functional connectivity with seed regions (the left precuneus and the left MIP) that showed a significant activation in the ([CB>SC] x [crossed > uncrossed]) contrast. In each individual, time-series of activity (first eigenvariate) were extracted from a 10mm sphere centered on the local maxima detected within 10 mm of the identified peaks in the second level analysis (SC>CB)x(Crossed>Uncrossed). New linear models were generated at the individual level, using three regressors. One regressor represented the condition (Crossed > Uncrossed). The second regressor was the activity extracted in the reference area. The third regressor represented the interaction of interest between the first (psychological) and the second (physiological) regressors. To build this regressor, the underlying neuronal activity was first estimated by a parametric empirical Bayes formulation, combined with the psychological factor and subsequently convolved with the hemodynamic response function (Gitelman et al., 2003). The design matrix also included movement parameters. A significant PPI indicated a change in the regression coefficients between any reported brain area and the reference region, related to the experimental condition (Crossed > Uncrossed). Next, individual summary statistic images obtained at the first level (fixed-effects) analysis were spatially smoothed (6-mm FWHM Gaussian kernel) and entered in a second-level (random-effects) analysis using a one-sample t-test contrasting CB>SC and SC>CB. Statistical inferences were conducted as for the main-effect analysis described above.

## Results

### Behavioral data

The slopes of each individual line (calculated from the z-scores of the mean percentages of “right hand first” responses) were submitted to an ANOVA with posture (uncrossed vs. crossed) as the within-subject factor and group (SC, CB) as the between-subject variable. Results showed: (1) a significant effect of posture [*F*(1, 17) = 6.52, *p* = .02, *η*^2^ = .28], the regression line for the uncrossed posture being steeper (*M* = .95 ± .01) than the regression line for the crossed posture (*M* = .58 ± .14); (2) a significant effect of group [*F*(1, 17) = 8.27, *p* = .01, *η*^2^ = .33], the CB (*M* = .97 ± .11) performing better (steeper regression) than the SC (*M* = .57 ± .09); and (3) a significant posture x group interaction [*F*(1, 17) = 6.75, *p* = .02, *η*^2^ = .28]. To further examine this interaction, paired samples t-tests compared hand positions in each group separately. In SC, participants’ performance was better in the uncrossed posture (*M* = .94 ± .02) than in the crossed posture (M = .20 ± .24), [*t*(10) = -3.04, *p* = .01]. In deep contrast, the CB group did not show any effect of posture [*t*(7) = 1.05, *p* = .33], the slope of the regression lines being similar in the uncrossed (*M* = .96 ± .004) and crossed postures (*M* = .97 ± .004). In SC, the Just Noticeable Difference (JND) was equal to 27 ms in the uncrossed position and 125 ms in the crossed posture. In the CB group, the JND was equal to 26 ms in both postures.

### fMRI data

We first tested whether our paradigm allowed us to observe the activation of the external remapping network in SC. Results revealed that the crossed condition, compared to the uncrossed posture, elicited brain responses in a large fronto-parietal network including the left superior parietal gyrus, the right posterior parietal cortex (PPC), the left precuneus, the left precentral gyrus, the left dorso-lateral prefrontal cortex, and the right middle temporal gyrus (see Fig. 1B and Table 2). The same contrast [crossed > uncrossed] performed in the CB group did not reveal any significant result. When the [crossed > uncrossed] contrast was directly compared between groups [SC vs CB], SC showed significantly more activity than the CB in the left precuneus, the left MIP, the left dorso-lateral prefrontal cortex and the right middle temporal gyrus (see Figure 1c and Table 2). CB did not show more activity than sighted for this contrast in any region.

**Figure 1.**
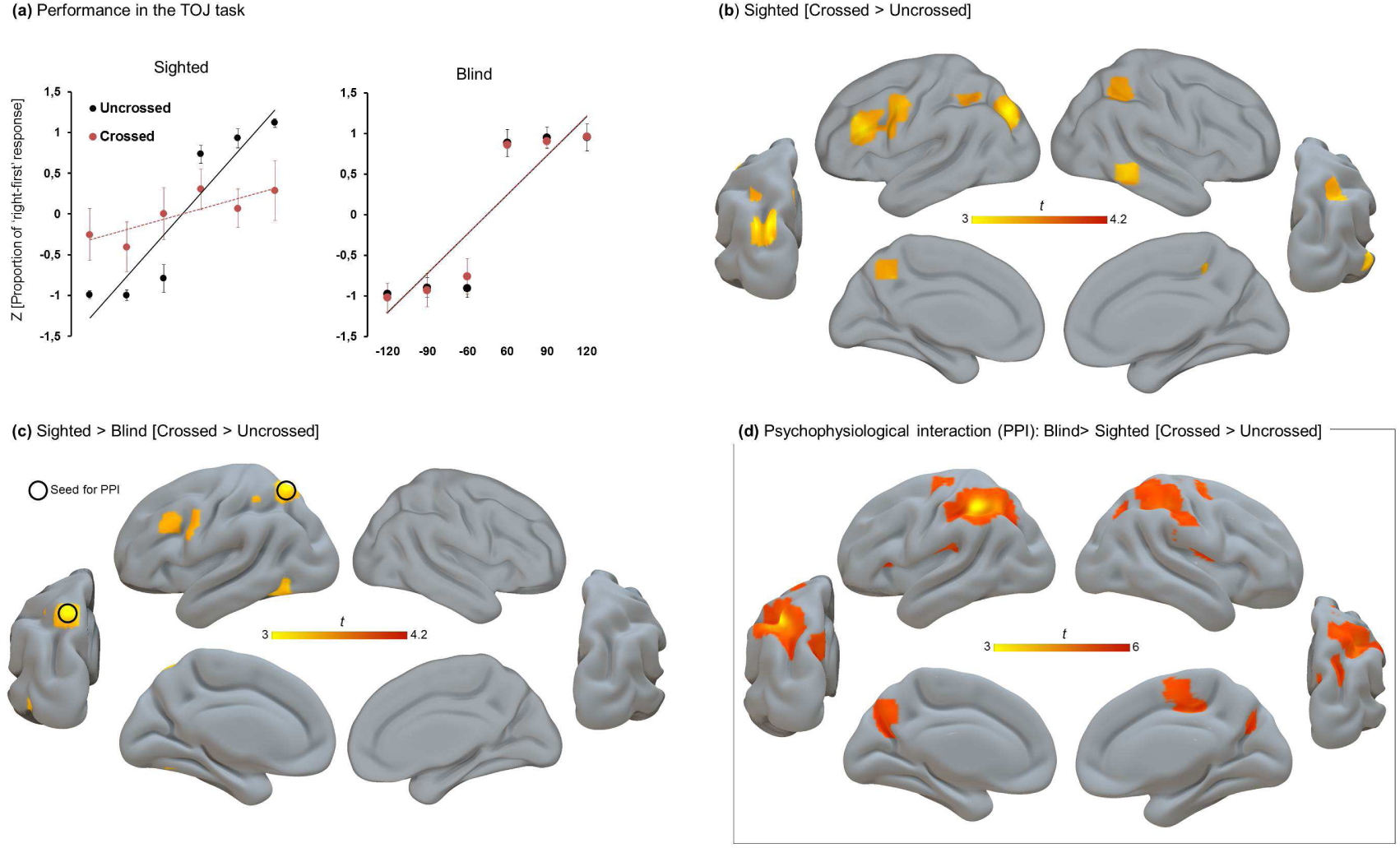
(A) Standardized z-score equivalents of the mean proportions of right-hand responses and best-fitting linear regression lines for the uncrossed (black lines) and crossed (red lines) postures for sighted and congenitally blind; (B) Results of the whole brain analyses probing brain activity obtained from the contrast testing which regions are specifically dedicated to the external remapping process in sighted ([Sighted] x [Crossed > Crossed]). There were no activations observed for this contrast in the blind group. (C) Regions selectively more active in the sighted group over the blind group in the crossed over the uncrossed posture ([Sighted > Blind] x [Crossed > Crossed]). (D) Functional connectivity changes. An increase of functional connectivity was observed between the left precuneus (seed encircled) and a bilateral fronto-parietal network when congenitally blind performed the TOJ task in the crossed over uncrossed posture. Whole brain maps are displayed at p<.001 uncorrected (k>15) for visualization purpose only (see methods for the assessment of statistical significance).

**Table 2.**
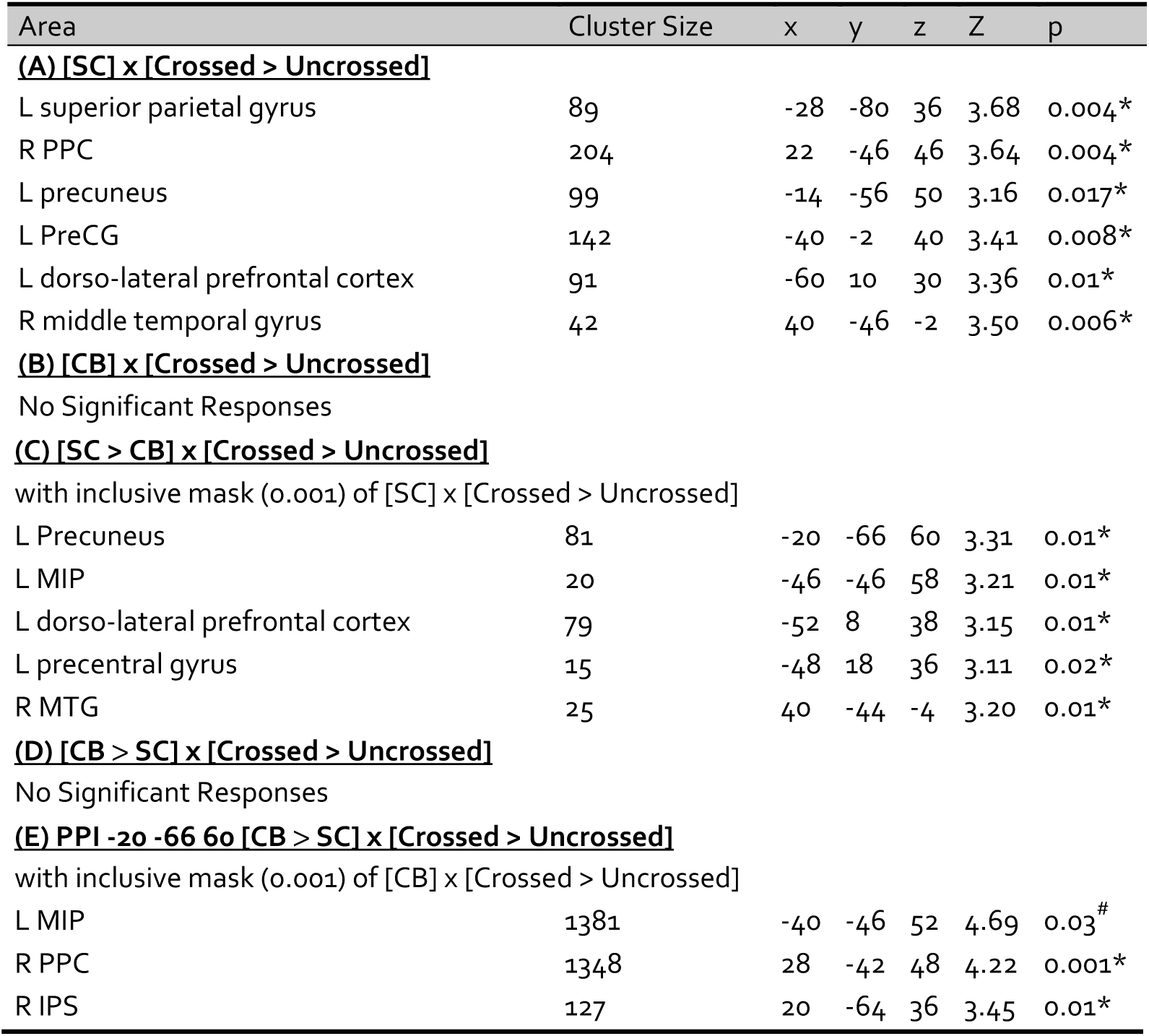
Functional results summarizing the main effect of groups for the different contrasts of interests

Psychophysiological interaction (PPI) analyses were computed to identify between-group differences in the functional connectivity maps of the regions involved in the automatic external remapping of touch identified in the sighted group. For these analyses, the left precuneus (-20, -66, 60 mm) was selected as seed region since it displayed the strongest between-group differences for the contrast [SC > CB] x [Crossed > Uncrossed] and also because this region was already reported in the literature as the neural basis of the external remapping of touch (Lloyd et al., 2003; Matsumoto et al., 2004; Azañón et al., 2010a; Takahashi et al., 2013; Wada et al., 2012). Interestingly, the results revealed that the seed regions showed stronger connectivity with and extended parietal network in CB compared to SC for the crossed over uncrossed posture (see Figure 1D).

Table 1. Brain activations significant (pcorr < .05 FWE) after correction over over the whole brain volume (^#^) or over small spherical volumes of interest (*). Cluster size represents the number of voxels in specific clusters when displayed at p(uncorr) < .001. SC: sighted controls, CB: congenitally blind, L: left, R: Right, MIP: medial intraparietal area, MTG: middle temporal gyrus, PPC: posterior parietal cortex; IPS: intraparietal sulcus.

## Discussion

We assessed the role visual experience plays in shaping the neural correlates of tactile localization. For this purpose, SC and CB participants were scanned while performing TOJ judgments with the hands uncrossed or crossed over the body midline. At a behavioral level, we observed that crossing the hands massively disrupted TOJ performance in SC but not in CB (see Figure 1A), replicating previous demonstration by Röder et al. (2004). While exploring the neurophysiological underpinning of this effect, we observed that the crossed condition, when compared to the uncrossed posture, elicited significantly more activity in the parietal and premotor areas in sighted, but not in blind participants. Our findings thus compellingly demonstrated that visual experience plays a crucial role in the development and/or engagement of a parieto-frontal network involved in this coordinate transformation process.

In sighted individuals, vision is a dominant sense for processing space due to the typically higher reliability, when compared to other senses, of the signal it provides for such a process. For instance, auditory or tactile information are typically remapped toward visual positions if inputs are spatially misaligned (Alais and Burr, 2004; Charbonneau et al., 2013); owls reared with prisms deviating their vision show permanent biases in auditory localization (Knudsen and Knudsen, 1989); and short-term adaptation to spatially conflicting visual and auditory stimuli biases auditory localization toward the visual source (Recanzone, 1998; Zwiers et al., 2003). Vision can even over-ride the proprioceptive sensation of a limb in space by displacing the position of a hidden arm toward a rubber one (Botvinick and Cohen, 1998). Actually, when we hear or feel something approaching or touching the body, we typically orient our vision toward this event and then use our motor system to guide appropriate action plans based on a precise location of the target in the external world (Goodale, 2011). As a result of their lack of visual experience, congenitally blind people have to rely exclusively on spatial information delivered by the remaining intact senses, such as hearing and touch. Thus, it seems likely that spatial perception in congenitally blind and in sighted people develops along different trajectories, and operates in a qualitatively different way in adulthood. Several studies have indeed pointed toward a reduced sense of external space in early blind individuals (Andersen et al., 1984; Bigelow, 1987; Dunlea, 1989; Millar, 1994; Ruggiero et al., 2012).

It has been shown that parietal and dorsal premotor regions play a crucial role in co-registering spatial information collected from various senses and frames of reference into a common coordinate system for the guidance of both eye and limb movements onto the external world (Graziano et al., 1994, 1997; Duhamel et al., 1998; Colby and Goldberg, 1999; Lloyd et al., 2003; Mullette-Gillman et al., 2005; Makin et al., 2007). For instance, it was shown that the position of the arm is represented in the premotor (Graziano, 1999) and parietal (Graziano, 2000) cortex of the monkey by means of a convergence of visual and proprioceptive cues onto the same neurons. More particularly, these regions are thought to be part of a network responsible for the remapping of skin-based touch representations located in somatosensory regions into external spatial coordinates (Lloyd et al., 2003; Matsumoto et al., 2004; Bolognini and Maravita, 2007; Zhaele et al., 2007; Azañón et al., 2010a; Longo et al., 2010; Takahashi et al., 2013; Wada et al., 2012). Accordingly, transiently disrupting the activity of the right posterior parietal cortex with Transcranial Magnetic Stimulation (TMS) selectively impairs the tactile remapping process but does not disrupt proprioceptive and somatosensory localization processes, highlighting the causal role of this region in remapping touch into external space (Azañón et al., 2010a).

When the hands are crossed, the conflict between external and anatomical representations of the hands increases the computational demands of the external remapping process which is typically observed in the “default” uncrossed posture (Melzack and Bromage, 1973; Bromage, 1974). Crossing the hands therefore triggers enhanced activity in the dorsal parieto-frontal network (see Figure 1B). In early blind people, the absence of a mandatory external remapping process prevents the increased recruitment of this neural network while crossing the hands. Therefore, by using blindness as a model system, we demonstrated that developmental vision plays a causal role in developing the computational architecture of parietal and dorsal premotor regions for transforming tactile coordinates from an initial skin-based representation to a representation that is defined by coordinates in external space.

Interestingly, it has recently been suggested that the integration of spatial information from different reference frames actually depends on the relative weight attributed to the internal and external coordinates (Azañón et al., 2010a; Badde et al., 2015; Badde and Heed, 2016). While integration seems mandatory in SC (Yamamoto and Kitazawa, 2001; Shore et al., 2002, Azañón et al., 2010b) the relative weight attributed to each coordinate system seems to be more dependent on tasks demands and instructions in CB (Heed and Röder, 2014; Heed et al., 2015; Crollen et al., 2017). Further studies should examine whether the external remapping network could therefore be active in CB while performing a task emphasizing external instructions. It is indeed possible that the external coordinate system is less automatically activated in CB than in SC but this does not mean that this system is not readily accessible when the task requires it (as, for example, when people perform an action directed toward the external world: Fiehler et al., 2009; Lingnau et al., 2014).

A recent study in the sighted demonstrated that the crossed-arms posture elicited stronger functional connectivity between the left IPS on the one hand and the right frontal gyrus and the left PPC on the other hand (Ora et al., 2016). By performing task-dependent functional connectivity analyses (Psychophysiological interactions), we demonstrate that blind individuals rely on enhanced integration between dorsal regions (Heine et al., 2015) while experiencing a conflict between body-centered and world-centered coordinates (see Figure 1D). This raises the intriguing possibility that changes in the connectivity pattern of the parietal cortex gates the activation, or not, of the external remapping process in congenitally blind people depending on task demands. Enhanced parieto-frontal connectivity in the crossed posture in the blind may therefore prevent the automatic remapping process from occurring in a task that does not necessitate such a computation (the TOJ task can be resolved by using pure skin-based coordinates). This could potentially explain the enhanced performance of the blind population in the crossed condition of the TOJ task (see Figure 1A).

In conclusion, we demonstrate that early visual deprivation alters the development of the brain network involved in the automatic multisensory integration of touch and proprioception into a common, external, spatial frame of reference. Moreover, the enhanced connectivity between dorsal regions in CB may provide a mechanistic framework to understand how blind people differently weight specific spatial coordinate systems depending on the task at play (Badde et al., 2015; Badde and Heed, 2016). These results have far-reaching implications for our understanding of how visual experience calibrates the development of brain networks dedicated to the spatial processing of touch.

## Acknowledgments

The authors are grateful to Giulia Dormal for her help in implementing the experimental design. This research and the authors were supported by the Canada Research Chair Program (FL), the Canadian Institutes of Health Research (FL), the Belgian National Funds for Scientific Research (VC), a WBI.World grant (VC), the European Union’s Horizon 2020 research and innovation programme under the Marie Sklodowska-Curie grant agreement No 700057 (VC) and the ‘MADVIS’ European Research Council starting grant (OC; ERC-StG 337573). O.C. is a research associate at the Belgian National Fund for Scientific Research. The authors declare no competing financial interest

## Author contributions

VC and OC designed research; VC, LL, and AB performed research; VC, OC, and MR analyzed data; VC and OC wrote the paper; FL provided laboratory resources for the optimal recruitment and testing of participants.

